# Frequent sgRNA-barcode Recombination in Single-cell Perturbation Assays

**DOI:** 10.1101/255638

**Authors:** Shiqi Xie, Anne Cooley, Daniel Armendarez, Pei Zhou, Gary C. Hon

## Abstract

Simultaneously detecting CRISPR-based perturbations and induced transcriptional changes in the same cell is a powerful approach to unraveling genome function. Several lentiviral approaches have been developed, some of which rely on the detection of distally located genetic barcodes as an indirect proxy of sgRNA identity. Since barcodes are often several kilobases from their corresponding sgRNAs, viral recombination-mediated swapping of barcodes and sgRNAs is feasible. Using a self-circularization-based sgRNA-barcode library preparation protocol, we estimate the recombination rate to be ~50% and we trace this phenomenon to the pooled viral packaging step. Recombination is random, and decreases the signal-to-noise ratio of the assay. Our results suggest that alternative approaches can increase the throughput and sensitivity of single-cell perturbation assays.

## Introduction

Recently, single-cell RNA sequencing (scRNA-seq) has been coupled with CRISPR-mediated perturbations, allowing functional assessment of genes (Perturb-seq, CRISP-seq, CROP-seq)[1–3] and enhancers (Mosaic-seq) [4] with a transcriptomic readout. All of these techniques deliver CRISPR components to cells through a lentiviral system, and each one has devised a unique strategy to detect sgRNAs through scRNA-Seq. Since the scRNA-seq strategies used are 3’-biased, most of these approaches insert a molecular barcode immediately before the poly(A) signal as an indirect proxy of sgRNA expression in each cell (**Figure. 1**). Therefore, the accuracy and sensitivity of these approaches rely on pre-identification of sgRNA-barcode relationships and unambiguous recovery of barcode information in every cell assayed.

**Figure 1.**
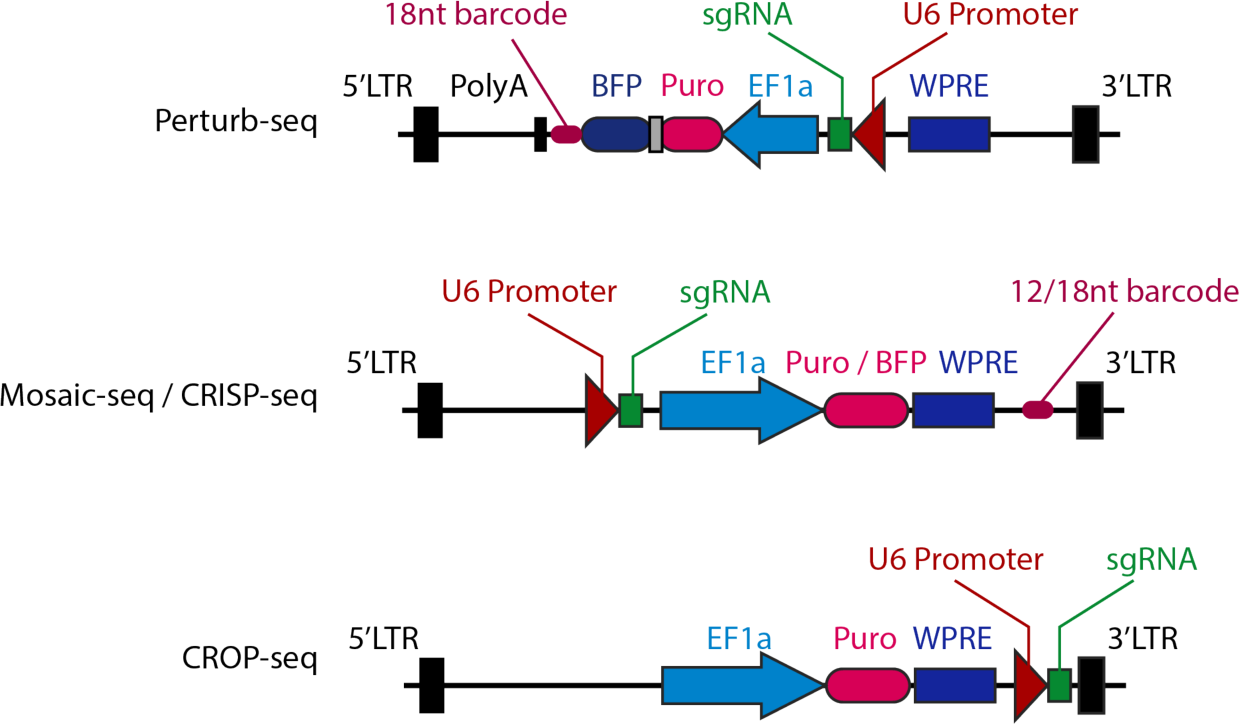
Vector structure of single-cell perturbation assays. The sgRNA barcode in Perturb-seq is part of the puromycin resistance gene / BFP transcript which is driven by core EF1a promoter (upper panel). Mosaic-seq and CRISP-seq share a similar design, in which the barcode is inserted immediately upstream of the lentiviral 3’LTR (middle panel). In CROP-seq, the whole sgRNA-expressing cassette is inserted between the WPRE and 3’LTR, allowing direct detection of sgRNA sequences.

However, barcoding could introduce noise due to lentiviral recombination. Two viral genomes are packaged into each lentiviral / retroviral particle[5], and are non-covalently linked[6]. During viral genome replication, the reverse transcriptase can switch from one template to another when it synthesizes a DNA provirus from a dimeric RNA genome, and this process happens most frequently at homologous regions[7–9]. The frequency of recombination depends on the distance between the two regions, which has been estimated to be 2% every kilobase[7,10]. Thus, when libraries of distinct sgRNA-barcode viruses are packaged together in single-cell perturbation assays, template switching could lead to barcode recombination that randomly shuffles sgRNA/barcode linkages. This event would interfere with the accurate detection of sgRNAs. A similar concern has also been raised recently on lentivirus-based genetic screening technologies[11].

## Results

To systematically measure the noise introduced by viral recombination during Mosaic-Seq, we individually cloned 21 unique sgRNAs into backbones with known barcode sequences (**Figure 2A and Methods**). We then monitored how pooling the samples at the transformation, viral packaging, or viral infection steps affected sgRNA-barcode recombination. To directly measure sgRNA-barcode pairs in each sample, we constructed deep sequencing libraries on plasmid pools and genomic DNA extracts. However, this problem is complicated by the large distance (~3-kb in Mosaic-Seq) separating each sgRNA to its barcode. Our strategy involves PCR amplification of ~3kb sgRNA-barcode amplicons followed by a self-circularization step, which reduces the sgRNA/barcode distance to ~400-bp (**Supplementary Methods**).

**Figure 2.**
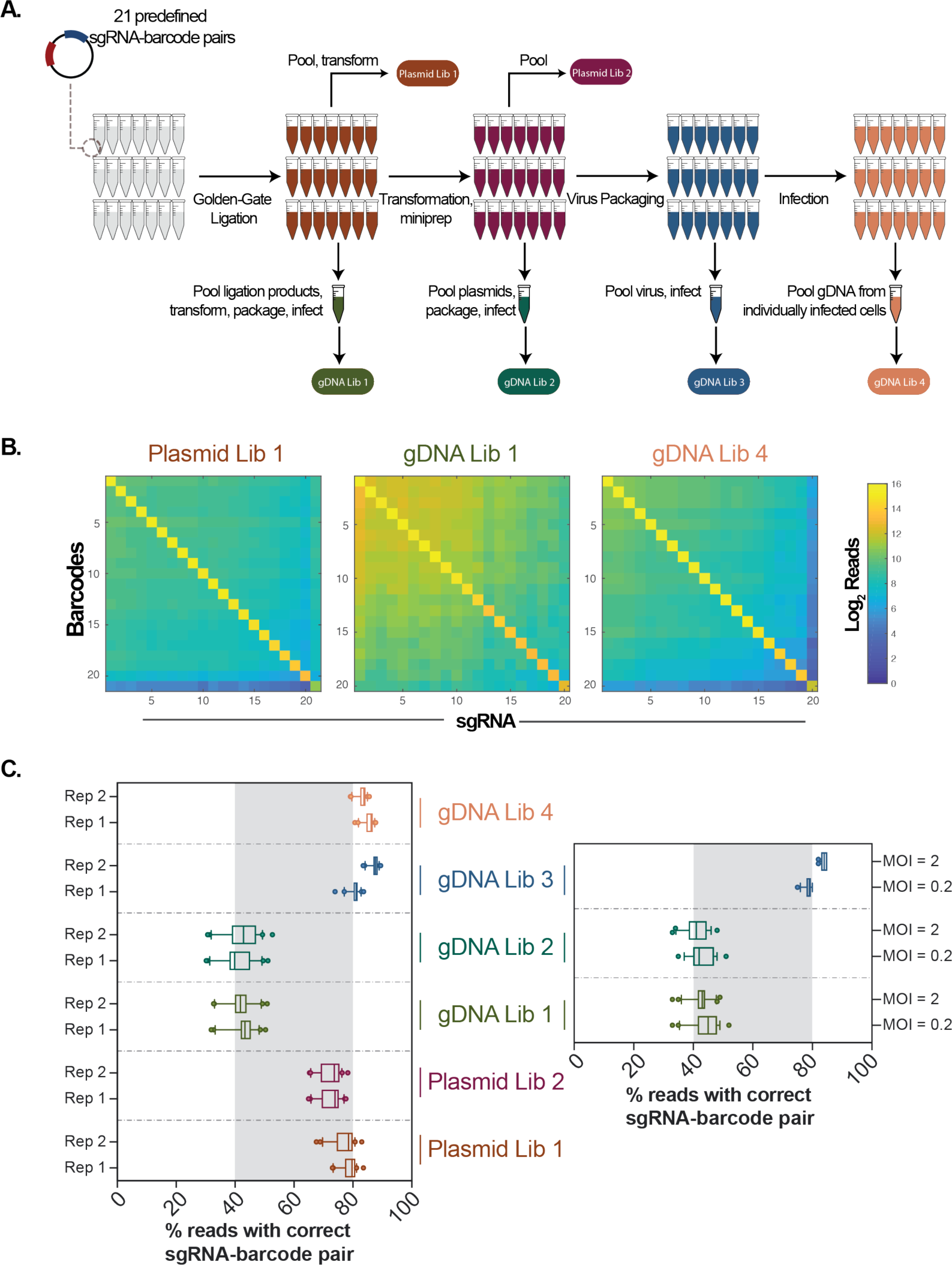
Barcode Shuffling during Multiplexed Mosaic-seq Library Preparation. (**A**) Schematic representation of the experiment design. 21 sgRNAs were inserted into 21 sgRNA backbones with distinct barcodes by Golden Gate assembly. Then the samples were pooled at different steps of the procedure and deep sequencing libraries were made from either plasmids or genomic DNA extracts of infected K562 cells (see methods). In total, we constructed two libraries from plasmids and four from the genomic DNA samples. (**B**) Read distribution of three representative libraries. For each barcode, we only observed one dominant sgRNA sequence, which is always the expected sgRNA. For the two libraries from genomic DNA samples, one sgRNA with ~100 reads dropped out. Therefore only 20 sgRNA-barcode pairs were used for heatmap plotting. (**C**) The percentage of reads for the most abundant sgRNA for each barcode were plotted by the boxplot. Whiskers represent the 10 and 90 percentiles and the dots represent outliers. Virus infection in the left panel were performed by using the same volume of virus stock (MOI varies from 1-2.4); right panel shows an independent experiment with high and low MOI, using the same virus in the left panel.

Since self-circularization is mediated by ligation, noise could be introduced by this method to assess recombination rates. To quantify this noise, we first examined Plasmid Library 2 (PL2), in which every sgRNA-barcode plasmid was constructed, transformed and extracted separately. We observe that 73.8% of reads (median) for each barcode is correctly linked to its known sgRNA pair, while the remaining 26.2% of reads are randomly linked to other sgRNAs (**Figure 2B, C**). This random collision rate correlates with the total abundance of each sgRNA in the library. As PL2 plasmids were independently processed, sgRNA-barcode recombination should be negligible. Therefore, 26.2% noise we observed is likely derived from our ligation-mediated method for detecting recombination.

Then, we examined recombination after pooled bacterial transformation (PL1). We also observed a median of 79.1% of reads exhibited correct sgRNA-barcode linkages, suggesting that pooled transformation does not significantly contribute to recombination in a library of sgRNA-barcode plasmids.

Next, we examined sgRNA-barcode pairs after viral integration into the human genome. At the four stages of Golden Gate ligation, transformation, viral packaging, and infection, samples were pooled, and sgRNA-barcode deep sequencing libraries were constructed on genomic DNA (**Figure 2A**). Two genomic DNA libraries in which sgRNA-barcode lentiviruses were individually packaged (GL3-4) maintain the correct sgRNA-barcode linkages (median of 83.6% and 84.4%, respectively) (**Figure 2C**), which is comparable to the plasmid libraries PL1-2.

In contrast, genomic DNA libraries in which plasmid libraries were pooled prior to viral packaging (GL1 and GL2), exhibited significant sgRNA-barcode recombination. The most abundant sgRNA of each barcode only occupies less than half of the reads (median of 42.4% and 41.3%, respectively), which is greater than a 50% loss compared to GL3-4. Recombination is random, and none of the incorrect sgRNA-barcode pairs are dominant over the expected pairs (**Figure 2B**). These results suggest that, using a strategy in which sgRNAs are separated from barocdes by several kilobases, recombination will be frequent if plasmid libraries are pooled prior to viral packaging.

To further test whether recombination depends on viral titer, we infected cells at high and low multiplicity of infection (MOI = 2 and MOI = 0.2). Based on Poisson statistics, >90% of antibiotic-selected cells are expected to be infected by exactly one virus at MOI = 0.2, which we hypothesize could reduce observed recombination rates compared to cells infected at high MOI. However, we observed no significant difference in recombination between the high and low MOI samples (**Figure 2C**), suggesting that the observed sgRNA-barcode shuffling is not due to recombination between multiple viruses infecting a single cell.

## Discussion

Our data confirms sgRNA-barcode recombination during pooled preparation of Mosaic-seq libraries. Recombination is random and accounts for ~50% of reads. While this noise is unlikely to create false positive hits, it does reduce the overall signal-to-noise of the assay, which we expect will decrease sensitivity. We postulate that similar recombination events will exist in other methods that rely on lentiviral/retroviral delivery systems that are coupled to indirect detection of DNA-based barcodes.

We observed that recombination only occurs when libraries are pooled before the virus production step, independent of the viral titer used during infection. This suggests that recombination predominantly occurs between two viral genomes packaged into the same virion, but not between distinct virus infecting the same cell. Thus, at low throughput, this problem can be overcome by constructing and packaging each virus separately. However, for large scale library preparation, the CROP-seq sgRNA plasmid is an improved solution[3]. In CROP-seq, the sgRNA cassette is inserted in proximal to the 3’LTR of the virus, which becomes part of the puromycin-resistance mRNA transcribed by EF1a promoter. Therefore, the sgRNA can be directly detected by scRNA-seq without the use of indirect barcodes. Moreover, CROP-seq dramatically simplifies the construction of large-scale sgRNA libraries since barcodes do not need to be constructed. By reducing sgRNA/barcode recombination, the sensitivity of single-cell perturbation assays could increase substantially. During preparation of this manuscript, similar observations have also been independently reported[12]. We believe that these improvements will significantly expand the application of single-cell perturbation assays, enabling the construction of large-scale libraries to systematically perturb and unravel transcriptional regulation from systems perspective.

## Methods

### Cell lines and culture

K562 cells were cultured in IMDM Medium plus 10% FBS and pen/strep at 37°C and 5% CO2. HEK293T cells were cultured in DMEM with 10% FBS and Pen/Strep. Both cells were acquired from ATCC.

### Plasmids

The lenti-sgRNA(MS2)-puro plasmid (Addgene ID: 73795) was used for sgRNA expression. The 12-bp barcode region flanking by a BsrGI and a EcoRI cutsite was inserted into this plasmid by using overlap PCR and Gibson assembly. For barcode insertion, a 108 bp oligo with 12bp random oligo sequence was synthesized and amplified by PCR in order to form double strand DNA. This fragment was then inserted into the linearized plasmid (cut with BsrGI and EcoRI) by Gibson assembly. After transformation, single clone was selected, and the barcode sequence of each clone was accessed by sanger sequencing. The insertion of sgRNA was performed by using BsmBI and T7 ligase, following the Golden Gate assembly protocol from Feng Zhang's lab[13]. To minimize the potential recombination, all the plasmids were transformed with Stellar Competent Cells (Clontech), and grown at 30°C.

### Virus Package, titration and infection

For virus packaging, 293T cells were seeded in a 6-cm dish (3X10^6^ cells) one day before transfection. The indicated viral plasmid(s) were co-transfected with lentiviral packaging plasmids pMD2.G and psPAX2 (Addgene ID 12259 and 12260) with 4:2:3 ratio by using linear polyethylenimine (PEI). Twelve hours after transfection, media was changed to fresh DMEM with 10% FBS plus Pen/Strep. Seventy-two hours after transfection, virus-containing media was collected, passed through a 45 µm filter, and aliquoted into 1.5ml tubes. Viruses were stored in −80 °C before infection or titration. Virus are then titrated and used for infection based on the methods described previously[4]. For infection of K562 cells, 2X10^5^ cells (in 500µl medium, with 8ng/µl polybrene) were used. After mixed with indicated amount of virus stock, the cells were centrifuged at 1000g, 37°C for 1 hour and then retracted to the incubator. The media was changed with fresh media containing 1µg/µl puromycin in the following day. The cells were selected for 7 days with media refreshment every two days and then collected for genomic DNA extraction and downstream library preparation.

### Construction of sequencing libraries

The method was described previously[4] with some modifications. Briefly, a 3kb amplicon with sgRNA and barcode on each end was amplified from plasmids or genomic DNA extracts. Then the fragment was self-circularized, and a second round of PCR was performed to yield a 400bp fragment with sgRNA and barcode adjacent to each other. See the **Supplementary Methods** for the detail of the protocol.

## Acknowledgements

We thank Vijay Ramani for his helpful insights into the sgRNA/barcode recombination problem, and we thank all the members in Hon lab for insightful discussion. This work is supported by the Cancer Prevention Research Institute of Texas (CPRIT) (RR140023, G.C.H.), NIH (DP2GM128203, G.C.H.), the Department of Defense (PR172060, G.C.H.), the Welch Foundation (I-1926-20170325, G.C.H.), and the Green Center for Reproductive Biology. S.X. is an American Heart Association fellow (16POST29910007). We acknowledge the BioHPC computational infrastructure at UT Southwestern for providing HPC and storage resources that have contributed to the research results reported within this paper. We also acknowledge UT Southwestern's McDermott Center for providing next-generation sequencing services for this work.

## Author Contributions

S.X. and G.C.H. conceived the study. S.X. and A.C. performed the experiments, analyzed the data. D.A. and P.Z. contributed to library preparation. S.X. and G.C.H prepared the manuscript.

